# Single-dose intranasal administration of AdCOVID elicits systemic and mucosal immunity against SARS-CoV-2 in mice

**DOI:** 10.1101/2020.10.10.331348

**Authors:** Rodney G King, Aaron Silva-Sanchez, Jessica N. Peel, Davide Botta, Selene Meza-Perez, S. Rameeza Allie, Michael D. Schultz, Mingyong Liu, John E. Bradley, Shihong Qiu, Guang Yang, Fen Zhou, Esther Zumaquero, Thomas S. Simpler, Betty Mousseau, John T. Killian, Brittany Dean, Qiao Shang, Jennifer L. Tipper, Christopher Risley, Kevin S. Harrod, Ray Feng, Young Lee, Bethlehem Shiberu, Vyjayanthi Krishnan, Isabelle Peguillet, Jianfeng Zhang, J. Todd Green, Troy D. Randall, Bertrand Georges, Frances E. Lund, Scot Roberts

## Abstract

The coronavirus disease 2019 (COVID-19) pandemic has highlighted the urgent need for effective preventive vaccination to reduce burden and spread of severe acute respiratory syndrome (SARS) coronavirus 2 (SARS-CoV-2) in humans. Intranasal vaccination is an attractive strategy to prevent COVID-19 as the nasal mucosa represents the first-line barrier to SARS-CoV-2 entry before viral spread to the lung. Although SARS-CoV-2 vaccine development is rapidly progressing, the current intramuscular vaccines are designed to elicit systemic immunity without conferring mucosal immunity. Here, we show that AdCOVID, an intranasal adenovirus type 5 (Ad5)-vectored vaccine encoding the receptor binding domain (RBD) of the SARS-CoV-2 spike protein, elicits a strong and focused immune response against RBD through the induction of mucosal IgA, serum neutralizing antibodies and CD4^+^ and CD8^+^ T cells with a Th1-like cytokine expression profile. Therefore, AdCOVID, which promotes concomitant systemic and local mucosal immunity, represents a promising COVID-19 vaccine candidate.

## Introduction

First reported in late 2019 in China (Zhu et al., 2020), infection caused severe acute respiratory syndrome (SARS) coronavirus 2 (SARS-CoV-2) has evolved into a global pandemic in just a few months. As of 5 October 2020, the World Health Organization estimates 34.8 million cases of coronavirus disease 2019 (COVID-19) worldwide, with 1,030,738 associated deaths (WHO. 2020a). Moreover, morbidity from SARS-CoV-2 infection can be severe, especially in high-risk groups (e.g., elderly, persons with chronic comorbidities such as hypertension, obesity, and diabetes) (CDC, 2020a). Early evidence from COVID-19 survivors and survivors of similar β-coronaviruses such as SARS and Middle East respiratory syndrome (MERS), suggests that COVID-19 survivors may suffer long-term sequela (e.g., inflammation of and damage to lungs and heart muscle) (Advisory Board, 2020). Taken together, the reach of infection and its impact on human health and well-being underscore the immediate need for safe and effective vaccines against SARS-CoV-2 to end this pandemic and prevent its return.

Despite the well-recognized role of mucosal immunity in prevention of disease (reviewed in Van Ginkel et al., 2000 and Holmgren et al., 2005), most of the COVID-19 vaccines in clinical testing are administered via intramuscular injection (Funk et al., 2020, WHO 2020b) – a route that elicits systemic immunity without inducing mucosal immune responses. The lack of mucosal immunity may limit the utility of intramuscularly administered COVID-19 vaccines, given that transmission of SARS-CoV-2 is primarily via respiratory droplets released by infected individuals in enclosed spaces (CDC, 2020b), with the nose and other portions of the respiratory mucosa being the primary routes of entry (Li et al., 2020). The nasal compartment shows particular susceptibility to SARS-CoV-2 infection due to abundant co-expression of the viral entry receptor (angiotensin-converting enzyme-2, ACE-2) and a required activating protease (TMPRSS2) in nasal goblet and ciliated cells (Sungnak et al., 2020). These cells are thought to be the most likely initial infection route for the virus and it is hypothesized that the nasal cavity serves as the initial reservoir for subsequent seeding of the virus to the lungs (Hou et al., 2020). The well-documented association of anosmia with COVID-19 further supports the nasal cavity as a principle reservoir of infection (Brann et al., 2020), and presence of high viral load in the nasal cavity may facilitate transmission of the virus. In contrast to intramuscular injection, mucosal vaccination via the intranasal route has the potential to confer sterilizing immunity in the respiratory tract (Hassan et al., 2020), reducing virus-induced disease and transmission of COVID-19.

Entry of SARS-CoV-2 into host cells depends on binding of the receptor-binding domain (RBD) of the spike protein to ACE-2, leading to fusion of the virus with the cell membrane (reviewed by Letko et al., 2020), and in human convalescent serum the majority of neutralizing antibodies are directed against RBD (Seydoux et al., 2020, Premkumar et al., 2020). While most clinically advanced SARS-CoV vaccine candidates deliver the trimeric spike ectodomain as the target antigen (Funk et al., 2020, WHO 2020b), subdomains of spike such as S1 and RBD represent alternative vaccine antigens for stimulating of a more focused immune response against these well-conserved domains, while limiting the induction non-functional antibodies and the risk of enhanced respiratory diseases (ERD).

Here, we report results of preclinical immunogenicity testing of AdCOVID, an intranasal replication-deficient Ad5-vectored vaccine candidate against COVID-19 that encodes the RBD from SARS-CoV-2 spike protein. Immunogenicity of the vaccine was assessed following a single administration in two strains of mice by measuring the induction of spike-specific antibody levels in sera and bronchoalveolar lavage (BAL) fluids. Functionality of these vaccine-elicited antibodies was measured in live virus neutralization assays. In addition to the induction of robust neutralizing antibody responses and mucosal IgA against SARS-CoV-2, the RBD vaccine candidate stimulated systemic and mucosal cell-mediated immune responses characterized by a T-helper 1 (Th1) type cytokine profile and through the induction of cytokine-producing CD4^+^ and CD8^+^ T cells, including lung-resident memory T (Trm) cells. These data, which demonstrate the immunogenicity of AdCOVID, support development of this candidate vaccine in response to the serious global health threat.

## Results

### Systemic and mucosal antibody responses induced by intranasal vaccine candidates expressing SARS-CoV-2 spike antigen subdomains

Three candidate COVID-19 vaccines were prepared using a replication-deficient E1- and E3-deleted Ad5 vector platform (Tang et al 2009). These Ad5 vectors were engineered to encode the human codon-optimized gene for full-length (FL) spike antigen (residues 1 to 1273), the S1 subdomain (residues 16 to 685) or RBD (residues 302 to 543) from the Wuhan-1 strain of SARS-CoV-2 (accession number QHD43416). Initial analysis of the FL vector indicated that antigen expression level and vector yield were significantly lower compared to the other two vaccine candidates, therefore, only the S1 and RBD vectors were evaluated in this study.

Immunogenicity of the S1 and RBD vectors was evaluated in both inbred C57BL/6J and outbred CD-1 mice. Mice received a single intranasal administration of the control vehicle or the Ad5 vectors tested at 3.35E+08 ifu (high dose), 6E+07 ifu (medium dose) or 6E+06 ifu (low dose) except for the high dose S1 vector that was tested in C57BL/6J mice at 6E+08 ifu. Following vaccine administration on day 0, serum and BAL samples were collected between days 7-28 (C57BL/6J) or days 7-21 (CD-1). IgG and IgA antibodies specific for SARS-CoV-2 spike were measured in serum or BAL samples using a spike cytometric bead array (CBA).

Systemic spike-specific IgG antibody responses in serum were detected in both strains of mice receiving a single intranasal administration of either the S1 or RBD vaccine **(Figure 1)**. The RBD vector induced a modestly larger serum spike-specific IgG response compared to the S1 vector – an effect that was more pronounced in the C57BL/6 mice despite the high dose RBD vector being half that of the high dose S1 vector. Moreover, intranasal administration of both the S1 vaccine and the RBD vaccine led to rapid elevation of spike-specific IgG **(Supplementary Figure 1)** in the BAL of both strains of mice.

**Figure 1:**
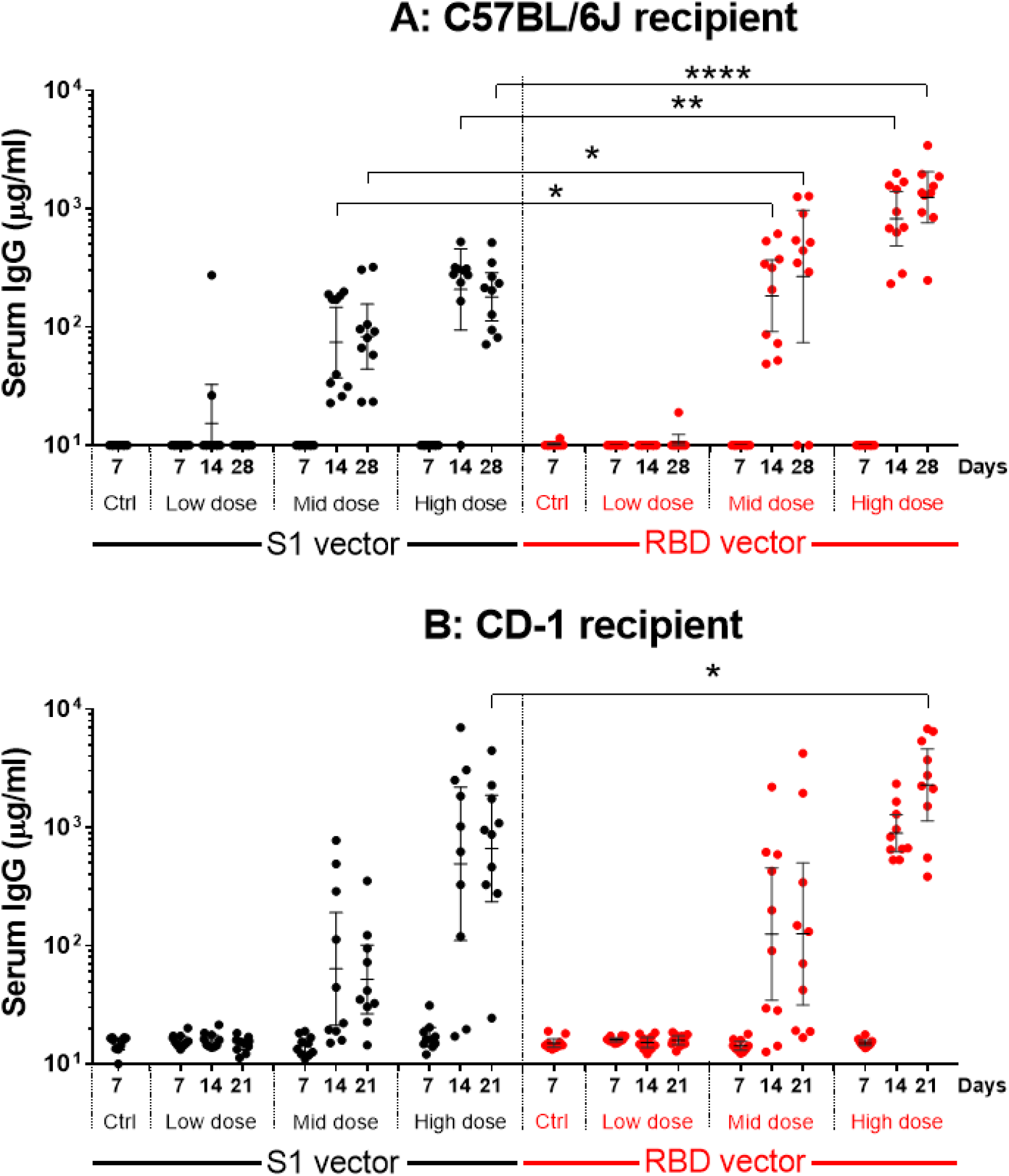
Spike-specific IgG responses in serum following single intranasal administration of S1 or RBD vectors. C57BL/6J (A) and CD-1 (B) mice received a single intranasal administration of vehicle (Ctrl), the S1 Ad5 vaccine (S1 vector) or RBD Ad5 vaccine (RBD vector) given at a low, middle (mid) or high dose as described in Material and Methods. Sera were collected between days 7-28 (panel A) or 7-12 (panel B) post-vaccination (n=10 animals/group/timepoint) and analyzed individually for quantification of spike-specific IgG. Results are expressed in µg/ml. Lines represent geometric mean response +/− 95% confidence interval. Statistical analysis was performed with a Mann-Whitney test: *, P < 0.05; **, P <0.01; ***, P < 0.001; ****, P < 0.0001.

The induction of mucosal immunity in BAL following a single intranasal dose of the S1 or RBD vaccine vectors was then assessed. Intranasal immunization is known to stimulate mucosal IgA antibodies, providing a first line of defense at the point of inoculation of respiratory pathogens (reviewed in Boyaka 2017). Pathogen-specific IgAs have previously been correlated with protection from respiratory infections such as influenza (Gould et al 2017, Belshe et al., 2000). As shown in Figure 2, both RBD and S1 vaccines induced a spike-specific IgA response at medium and high doses of vaccine. The reason for the markedly greater induction if IgA in CD-1 mice is not known but may be related to mouse strain-specific predisposition to produce innate and vaccine-induced mucosal IgA (Franssen et al., 2015).

**Figure 2:**
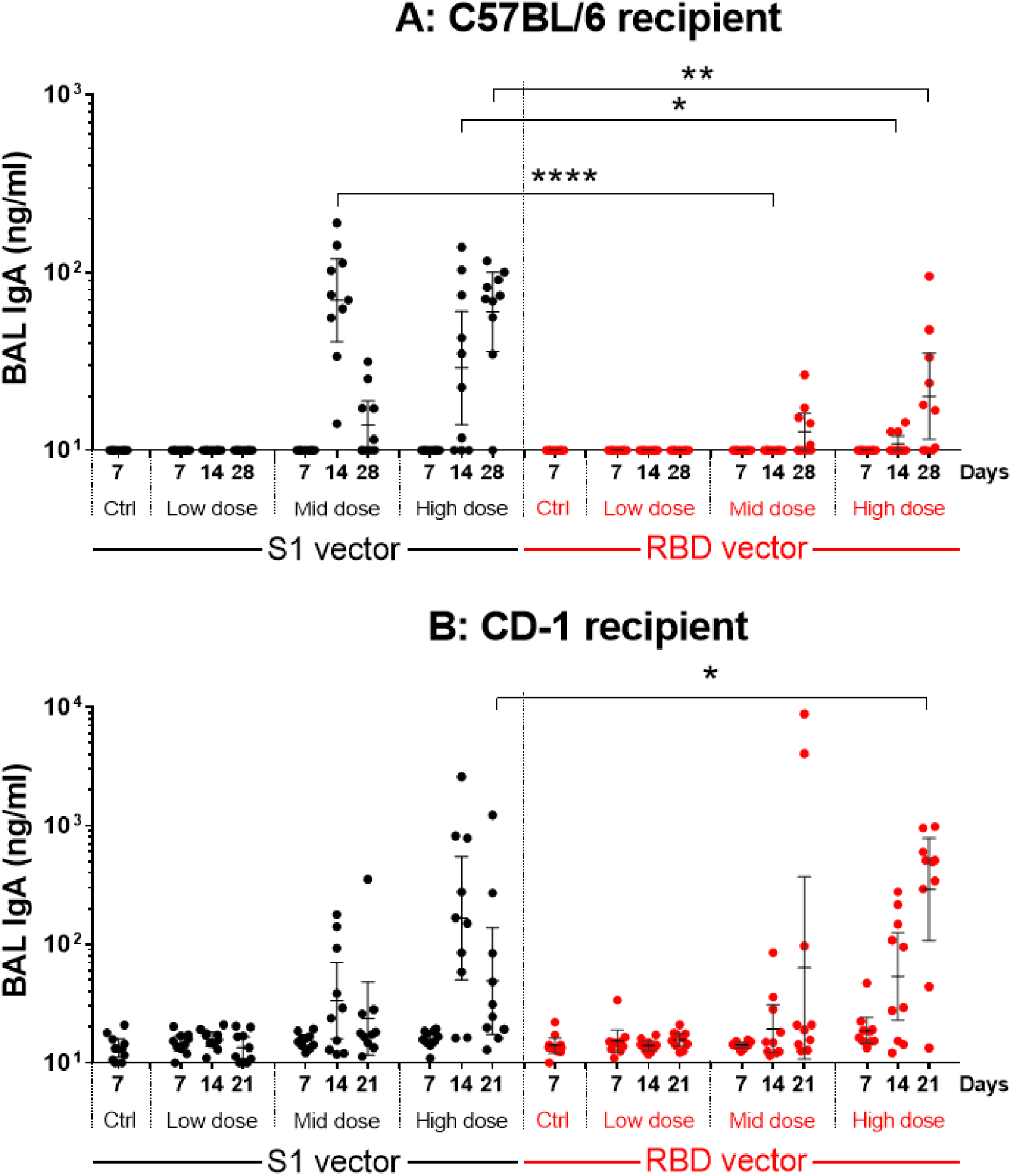
Spike-specific IgA responses in BALs following single intranasal administration of S1 and RBD vectors. C57BL/6J (A) and CD-1 (B) mice received a single intranasal administration of vehicle (Ctrl), the S1 Ad5 vaccine (S1 vector) or RBD Ad5 vaccine (RBD vector) given at a low, middle (mid) or high dose as described in Material and Methods. BAL samples were collected at the indicated timepoints (n=10 animals/group/timepoint) and analyzed individually for the quantification of Spike-specific IgA. Results are expressed in ng/ml. Lines represent geometric mean response +/− 95% confidence interval. Statistical analysis was performed with a Mann-Whitney test: *, P < 0.05; **, P <0.01; ***, P < 0.001; ****, P < 0.0001.

Vaccine-elicited neutralizing antibody responses against SARS-CoV-2 were measured in a focus reduction neutralization test (FRNT) using the wild-type SARS-CoV-2 isolate USA-WA1/2020. The analysis included S1 and RBD samples from C57BL/6J mice 4-weeks after vaccination with either medium or high-dose vaccine, and RBD samples from CD-1 mice 3-weeks after vaccination with the high vaccine dose (**Figure 3**). RBD elicited significantly greater neutralizing titers compared to S1 under all conditions evaluated. At the highest dose, intranasal vaccination with RBD induced neutralizing antibody responses above background in 10 of 10 C57BL/6J mice and 8 of 10 CD-1 mice with a median titer of 563 and 431, respectively **(Figure 3A)**. The level of the neutralizing antibody response was well-correlated with magnitude of the spike-specific serum IgG response measured in individual animals **(Figure 3B and 3C)**, indicating that robust antibody responses to RBD were associated with generation of potentially protective neutralizing antibodies.

**Figure 3:**
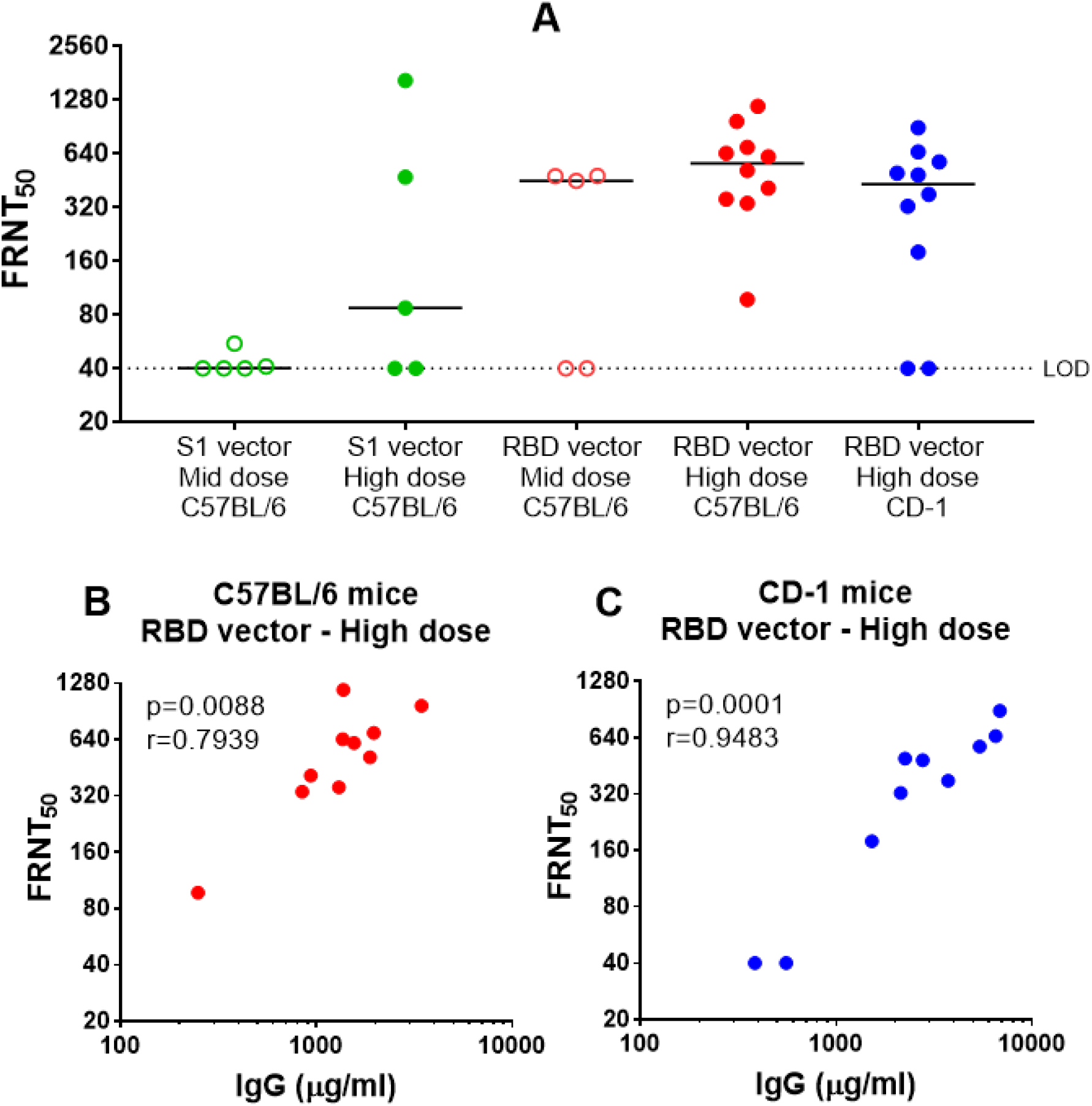
SARS-CoV-2 neutralizing antibody responses in serum following single intranasal administration of the S1 and RBD vectors. **(A)** Neutralizing antibody response by C57BL/6J or CD-1 mice vaccinated 28 days earlier with the mid or high dose of the S1 or RBD vector as indicated. Results are expressed as the reciprocal of the dilution of serum samples required to achieve 50% neutralization (FRNT_50_) of wild-type SARS-CoV-2 infection of permissive Vero E6 cells. Line represent the group median value. **(B-C)** Correlation between neutralizing antibody response and Spike-specific IgG response in serum of vaccinated animals. Correlation analysis was performed with a two-tailed Spearman test.

### Recruitment of innate and adaptive immune cells in the respiratory tract following intranasal RBD vector administration

Given the markedly greater neutralization titers obtained with the RBD vaccine candidate compared to the S1 candidate, the RBD vaccine was further evaluated with respect to cellular immunity. Flow cytometric analysis of immune cells was performed in BAL, lung, mediastinal lymph node (medLN) and spleen samples following intranasal administration of the RBD vector in C57BL/6J mice. Consistent with the hypothesis that mucosal administration of the vaccine would induce innate and adaptive pulmonary immune responses, rapid recruitment of immune cells into the lung was observed following intranasal vaccination with the RBD vector (**Figure 4**). Indicative of an early innate immune activation, significant increases in the number of dendritic cells, macrophages and natural killer cells were observed, which peaked at day 7 post-vaccination. Rapid recruitment of adaptive immune cells including T follicular helper cells (Tfh) and B cells to the lung were also observed peaking between days 7-28. Similar trends were observed in the medLN (**Supplementary Figure 2**) and BAL (**Supplementary Figure 3**) but were less obvious in the spleen (**Supplementary Figure 4**).

**Figure 4:**
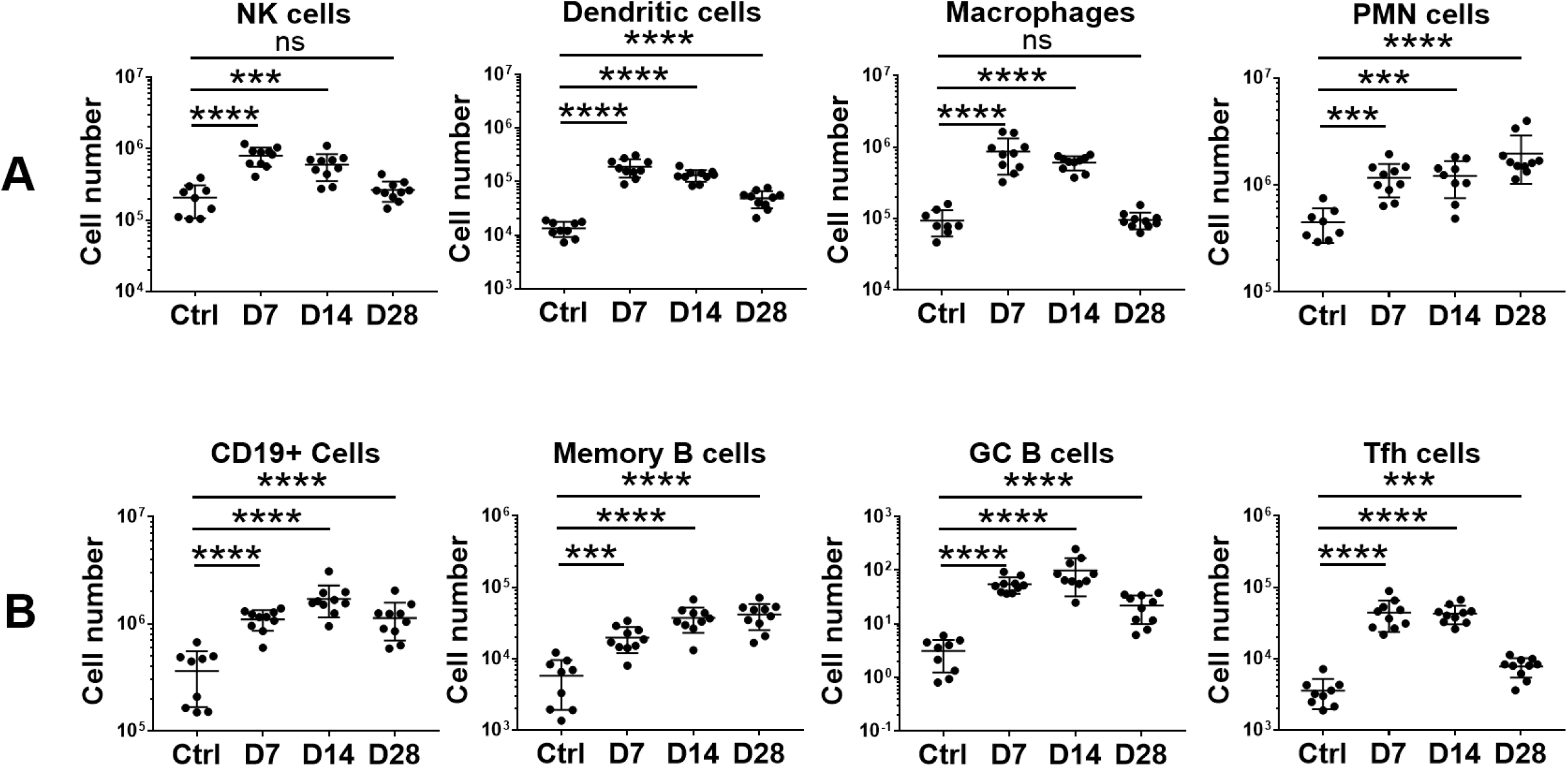
Flow cytometry analysis of immune cells in lungs from C57BL/6J mice following single intranasal high dose of RBD vector (AdCOVID). C57BL/6J mice were given a single intranasal administration of vehicle (Ctrl) or the high dose RBD vector (AdCOVID) as described in Material and Methods. Lung cells were isolated from the vaccinated mice at the timepoints indicated (10 mice/timepoint) and analyzed individually by flow cytometry using markers of (A) innate immune cells or (B) B and Tfh cells as described in Material and Methods. Results are expressed as cell number. Lines represent mean response +/− SD. Statistical analysis was performed with a Mann-Whitney test: *, P < 0.05; **, P <0.01; ***, P < 0.001; ****, P < 0.0001.

### Mucosal and systemic antigen-specific CD4+ and CD8+ T cell responses

Emerging data support a role for T cell responses in COVID-19 immunity independent of antibody responses (Sekine et al., 2020). In addition, animal models have shown that CD4^+^ and CD8^+^ T cells that reside in the respiratory tract are important for local protection immediately after viral infection (Pizzolla et al., 2017, Laidlaw et al., 2014, Takamura et al., 2017).

To assess vaccine-induced SARS-CoV-2-specific T cell responses, RBD Ad5 vector (also referred to AdCOVID) was administered intranasally to outbred CD-1 mice at a single high dose. Control animals received vehicle alone administered intranasally. RBD-specific T cell cytokine responses in lung and spleen samples were assessed following *ex vivo* re-stimulation with a pool of 54 peptides (15a.a. long, 11a.a. overlap) covering the SARS-CoV-2 RBD residues 319-541.

As measured by ELISpot, a high frequency of IFN-γ-producing RBD-specific T cells were detected in the lung at 10- and 14-days post-vaccination, reaching a mean response of 915 and 706 spot forming cells per million input cells respectively **(Figure 5)**. IFN-γ producing RBD-specific T cells were also detected by ELISpot in the spleen, albeit at lower frequency compared to lung. This suggests that functional effector T cells primed in response to mucosally-delivered antigens can migrate to peripheral lymphoid tissues.

**Figure 5:**
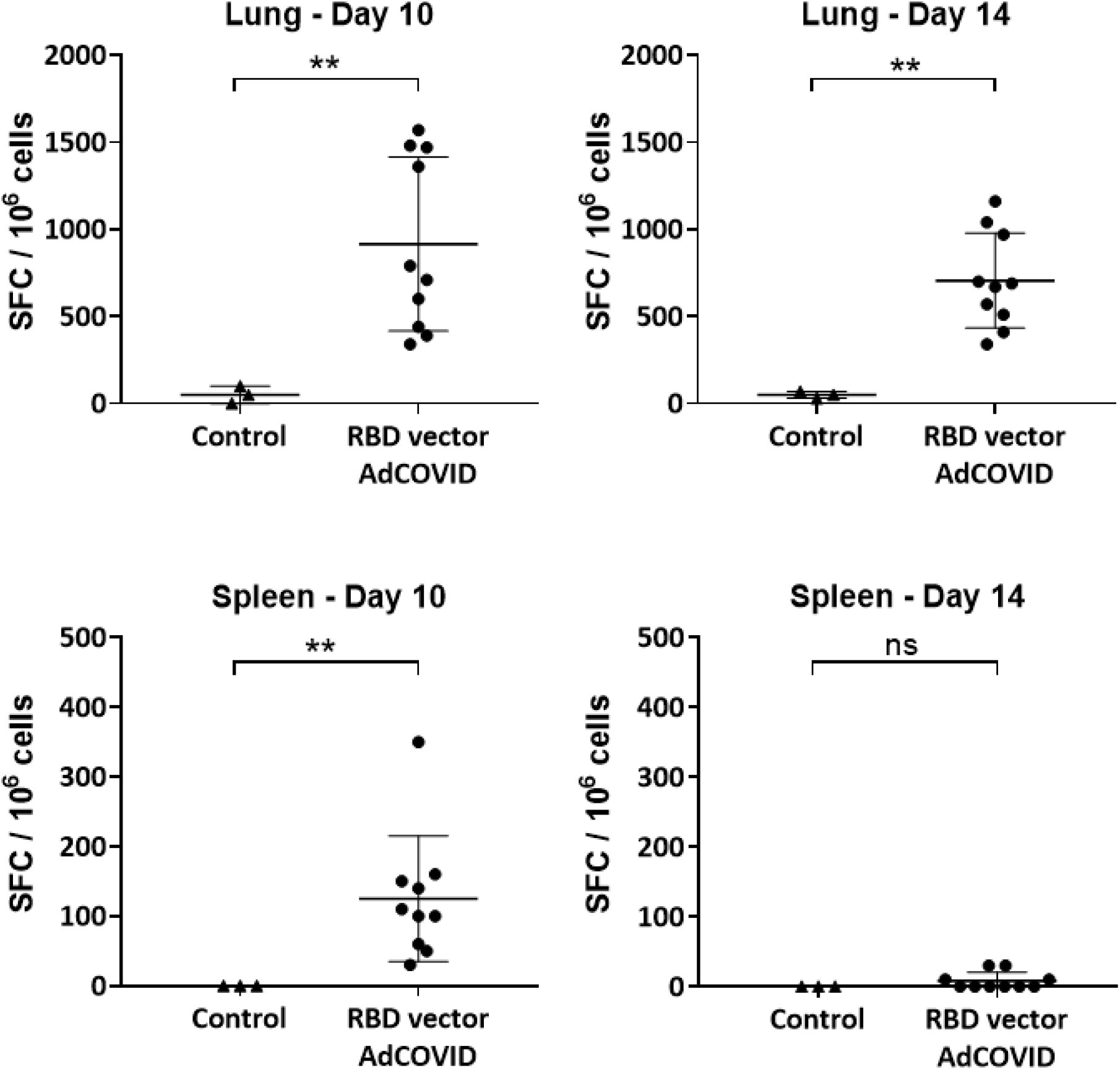
IFN-γ ELISpot responses of T cells in lungs and spleen of CD-1 mice following single intranasal high dose vaccination with RBD vector (AdCOVID). C57BL/6J mice were given a single intranasal administration of vehicle (Ctrl) or the high dose RBD vector (AdCOVID) as described in Material and Methods. Lung and spleen cells were isolated on days 10 and 14 following immunization, re-stimulated with an RBD peptide pool and analyzed by an IFN-γ ELISpot assay. Results are expressed as Spot Forming Cells (SFC) per million input cells. Lines represent mean response +/− SD. Statistical analysis was performed with a Mann-Whitney test: *, P < 0.05; **, P <0.01; ***, P < 0.001; ****, P < 0.0001.

To further characterize the RBD-specific CD4^+^ and CD8^+^ T cell responses to mucosal vaccination, intracellular cytokine staining was performed on lung and spleen cells from vaccinated mice. Consistent with ELISpot data, we observed a significant induction of IFN-γ- or TNF-α-producing T cells in the lung and spleen (**Figure 6**) of vaccinated animals. These included both CD11a^+^CD4^+^ and CD11a^+^CD8^+^ T cells, although response was strongly biased toward the induction of CD8^+^ T cells, especially in lung samples. Expression of integrin CD11a, which is only upregulated in recently activated T cells and is required for optimal vascular adhesion in the tissue and retention within the respiratory tract (Thatte et al.. 2003), supports the hypothesis that these cells were recently recruited to the lung. To assess whether these cells are resident memory T cells (Trm), the expression of the Trm markers CD103 and CD69 (Takamura, 2017) on pulmonary CD4^+^ and CD8^+^ cells was determined. Consistent with the intranasal administration route, induction of lung RDB-specific CD4^+^ and CD8^+^ Trm expressing either IFN-γ, TNF-α or both cytokines were observed (**Figure 7**).

**Figure 6:**
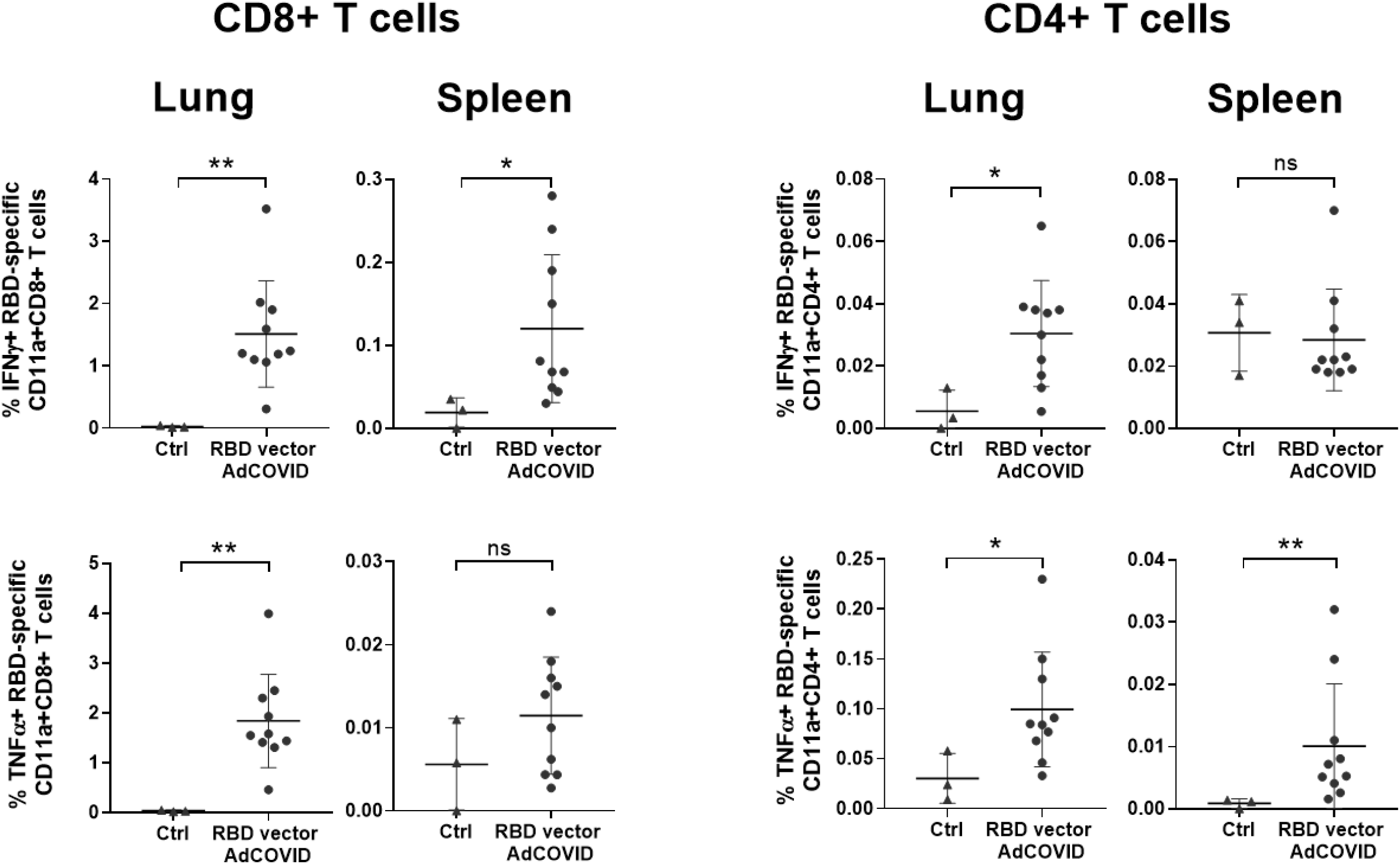
Intracellular cytokine production by pulmonary and spleen T cells 14 days after intranasal administration of RBD vector (AdCOVID). CD-1 mice were given a single intranasal administration of vehicle (Ctrl) or high dose RBD vector (AdCOVID) as described in Material and Methods. Lung cells (n= 10 mice/vaccine, 3 mice/control) were isolated on day 14, re-stimulated with the RBD peptide pool for 5 hrs and analyzed by flow cytometry. Results are expressed as the % of IFN-γ or TNF-α expressing CD11a^+^ CD4^+^/CD8^+^ T cells for individual mice. Different Y-axis scales are used across the graphics. Lines represent mean response +/− SD. Statistical analysis was performed with a Mann-Whitney test: *, P < 0.05; **, P <0.01; ***, P < 0.001; ****, P < 0.0001.

**Figure 7:**
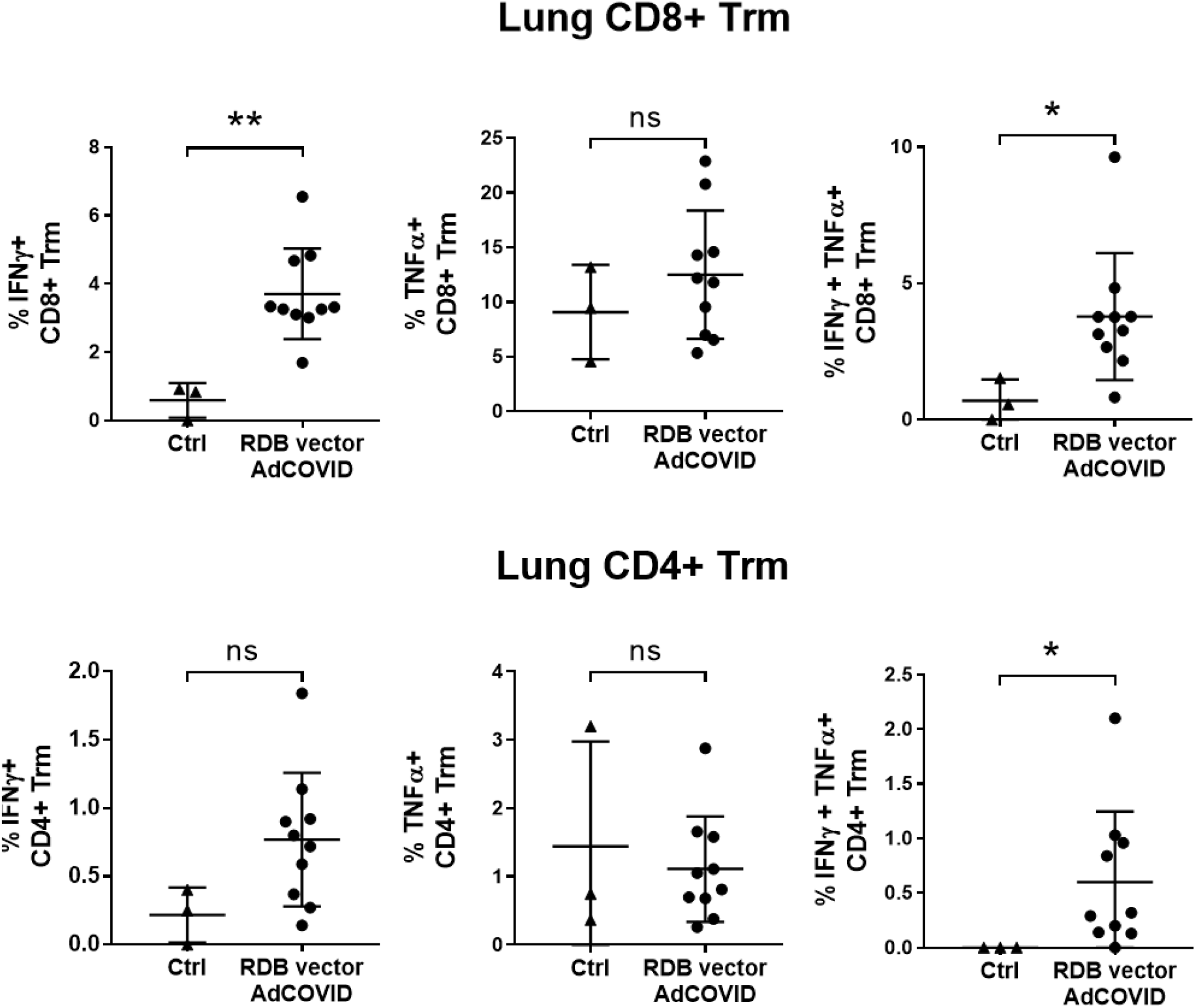
Intracellular cytokine production by lung resident memory T cells at 14 days after single intranasal administration with RBD vector (AdCOVID). CD-1 mice were given a single intranasal administration of vehicle (Ctrl) or high dose RBD vector (AdCOVID) as described in Material and Methods. Lung cells (n= 10 mice/vaccine, 3 mice/control) were isolated at day 14, re-stimulated with the RBD peptide pool for 5 hrs and analyzed by flow cytometry to identify CD69^+^CD103^+^ resident memory T cells (Trm). Results are expressed as the % of IFN-γ+, TNF-α+ or double positive IFN-γ+/TNF-α+ expressing CD4+ or CD8+ Trm cells for individual mice. Lines presented as the mean response +/− SD for the groups. Statistical analysis was performed with a Mann-Whitney test: *, P < 0.05; **, P <0.01; ***, P < 0.001; ****, P < 0.0001.

Our data showed that intranasal administration of RBD vaccine candidate induced T cells competent to produce IFN-γ and TNF-α – cytokines that are associated with Th-1 biased cellular response. In addition, the vaccine elicited high frequencies of antigen-specific CD8^+^ T cells that generally correlate with an IFN-regulated T cell response that is important for control of viral infection. To further assess the cytokine producing potential of the T cells from vaccinated mice, Splenic T cells were stimulated with RBD peptides for 48 hours and then used cytokine bead arrays to measure cytokine levels in the supernatant (**Figure 8**). As expected, induction of IFN-γ and TNF-α by the T cells was observed. Moreover, T cells from the vaccinated animals produced moderate levels of IL-10 compared to T cells from the vehicle control treated mice. Importantly, Interleukin (IL)-4, IL-5 and IL-13 and IL-17a levels in the supernatant from re-stimulated cells derived from the vaccinated mice were equivalent to that seen in cultures containing peptide-stimulated cells from the vehicle control animals. These data therefore indicated that intranasal administration of the RBD vaccine did not initiate a potentially deleterious Th2 response but rather induced the expected antiviral T cell responses.

**Figure 8:**
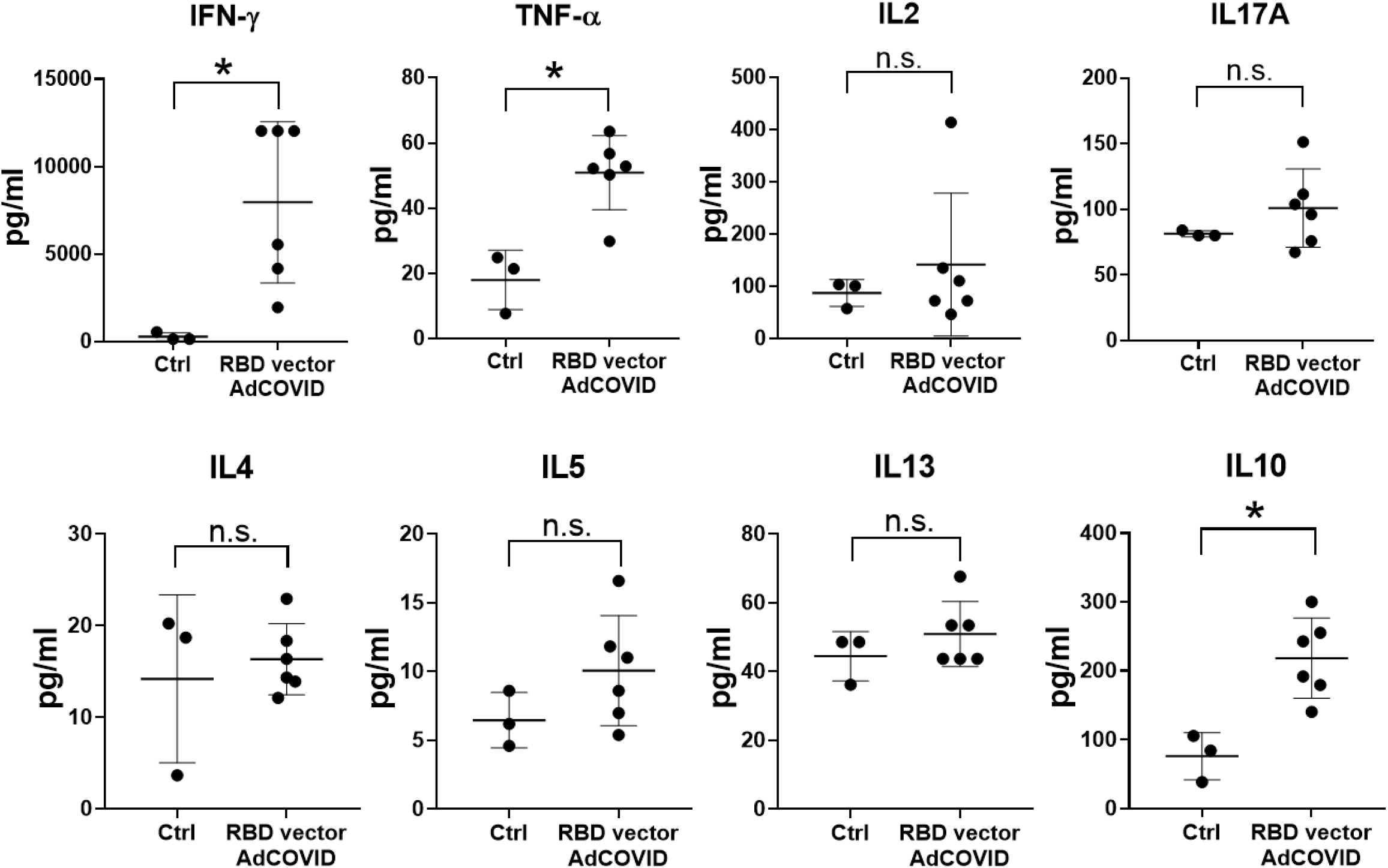
Secreted cytokine production by splenic T cells 10 days after single intranasal administration with RBD vector (AdCOVID). CD-1 mice were given a single intranasal administration of vehicle (Ctrl) or high dose RBD vector (AdCOVID) as described in Material and Methods. Spleen cells (n= 10 mice/vaccine, 3 mice/control) were isolated at day 10 and re-stimulated with the RBD peptide pool for 48 hrs. Secreted cytokines were detected in the supernatant using a cytokine multiplex assay. Results are expressed in pg/ml. Lines represent mean response +/− SD. Statistical analysis was performed with a Mann-Whitney test: *, P < 0.05; **, P <0.01; ***, P < 0.001; ****, P < 0.0001

Taken altogether, the data show that intranasal administration of the replication-deficient Ad5 vector expressing SARS-CoV-2 spike RBD generates humoral and cellular immune responses in both systemic and mucosal sites, particularly within the lungs, which represents a major site for infection and clinical disease.

### Discussion

We demonstrate that AdCOVID, an intranasal adenovirus-vectored vaccine encoding the RBD of the SARS-CoV-2 spike protein is highly immunogenic in both inbred and outbred mice and elicits both robust systemic and local antibody and T cell responses. Following a single intranasal vaccination, AdCOVID elicited a strong and focused immune response against SARS-CoV-2 spike antigen through the induction of functional serum antibodies that neutralize wild-type SARS-CoV-2 infection as well as mucosal IgA and polyfunctional CD4^+^ and CD8^+^ T cell responses in the respiratory tract. Cell-mediated responses induced by AdCOVID were biased toward an antiviral response as demonstrated by the high rates of antigen-specific CD8^+^ T cells and a cytokine expression profile including IFN-γ and TNF-α. Establishment of a resident memory CD8+ T cell population in the lungs complements robust induction of mucosal IgA antibody against the spike protein, an important addition to the overall immune response to AdCOVID.

An interesting observation from our studies was the lower neutralizing antibody titers in animals receiving the S1 vaccine. We hypothesize that the RBD vector better focused the immune response on the critical neutralizing epitopes in the spike protein, and similar results were obtained during the development of vaccines for SARS (Ravichandran et al., 2020). Regardless, the selection of an Ad5-vector expressing RBD offers two advantages over other forms of the spike antigen used in COVID-19 vaccine candidates currently in clinical development. First, and in agreement with our data, RBD-based vaccines were previously shown to promote equivalent or better antibody responses than the full length or S1/S2 ectodomain of spike from SARS-CoV-2 including higher affinity interaction compared to the S1 domain (Ravichandran et al., 2020, Walsh et al., 2020). Second, focusing the immune response on the RBD domain could decrease the production of potentially pathogenic non-neutralizing antibodies, which can contribute to enhanced respiratory diseases (ERD).

To date, the COVID-19 vaccine candidates that have advanced to Phase 3 clinical trials have been based on the intramuscular injection of the vaccine candidate, and there is no widespread expectation that the current intramuscular vaccine candidates will provide sterilizing immunity in the nasal cavity even if they prove effective in reducing mild to moderate COVID-19 disease. In preclinical and clinical studies, these candidates have demonstrated stimulation of serum neutralizing antibodies, and peripheral T cell responses. However, a fundamental limitation of these approaches is that they are not designed to elicit mucosal immunity. In a SARS-CoV-2 challenge model in rhesus macaques, a single intramuscular administration of an adenovectored vaccine candidate was shown to significantly reduce viral load in the bronchoalveolar lavage fluid and lower respiratory tract tissue but the level of viral replication in the nasal cavity was unaffected (van Doremalen at al., 2020). Consistent with that report, several studies assessing other COVID-19 vaccine candidates have shown that the intranasal route, as opposed to the intramuscular route, stimulated local mucosal immune responses in addition systemic neutralizing antibody and T cell responses and achieved sterilizing immunity (Hassan et al., 2020; Wu et al., 2020). Moreover, several studies have demonstrated the ability of intranasally administered vaccines to block transmission of influenza between infected and naïve cage-mates (Lowen et al., 2009; Price et al., 2011), Nasal administration of replication-deficient human adenovirus serotype 5 (Ad5)-vectored vaccines mimics a route of natural infection and is known to stimulate strong humoral and cellular immunity, both systemically and mucosally (Croyle et al., 2008). Moreover, intranasal Ad5-vectored vaccine have been shown to provide effective protection against respiratory pathogens (Zhang et al., 2011; Krishnan et al., 2015), and the intranasal route of administration has been demonstrated to bypass preexisting immunity to the vector (Croyle et al., 2008). Another advantage of intranasal AdCOVID vaccine, compared to other vaccines in development, is its ease of administration and delivery to patients, as intranasal administration is non-invasive and obviates the need for needles which may increase vaccine uptake (reviewed in Yusuf et al., 2017).

In summary, there is a clear unmet need for a vaccine to prevent SARS-CoV-2 infection and its transmission. In these experiments, AdCOVID induced systemic and mucosal immune responses within days following single-dose immunization. In the context of a pandemic, AdCOVID has two compelling advantages. First, non-invasive intranasal administration makes it particularly well-suited for wide-spread vaccinations of large cohorts. Second, AdCOVID may offer the ability to control SARS-CoV-2 infection within the upper and lower respiratory tract. This has the advantages of potentially preventing infection at the site of virus entry, reducing the risk of significant respiratory disease and decreasing the likelihood of subsequent virus transmission in vaccinated individuals. Collectively, these findings support further development of AdCOVID for the prevention of SARS-CoV-2 infections and its transmission.

## Acknowledgements

Altimmune Inc. and the Barbara Ingalls Shook Foundation (to FEL) provided funding for the work reported in this article. Support for the development and validation of SARS-CoV-2 spike cytometric bead arrays was provided by U19 3U19AI142737 (to FEL, RGK, JTG, TDR and Dr. John F. Kearney). We thank Dr. Kearney for his helpful suggestions and support of this project. We thank Ms. Uma Mudurunu and the many UAB Animal Resource Program staff who provided care for the vaccinated animals during the university shutdown due to COVID-19.

## Author Contributions

RGK, JTG, TDR, BG, FEL and SR participated in study design and data interpretation. RGK, AS-S, JNP, DB, SM-P, SRA, MDS, ML, JLT and KSH participated in data analysis. RGK, AS-S, JNP, DB, SM-P, SRA, MDS, ML, JEB, SQ, GY, FZ, EZ, TSS, BM, JTK, BD, QS, JLT and CR participated in reagent development and data collection. BG, SR, RF, YL, BS, VK, IP and JZ design and/or produced and tested the vaccine candidates. BG and FEL wrote the initial draft; the other authors reviewed and provided editorial input into subsequent drafts. All authors had full access to all the data in the study and take responsibility for integrity of the data and the accuracy of the data analysis.

## Potential Conflict of Interest

The authors located at University of Alabama at Birmingham declare no potential conflicts of interest. BG, SR, RF, YL, BS, VK, IP and JZ are employees of Altimmune Inc. and received stock options and compensation as part of their employment.

## Material and Methods

### Ethics statement and Mice

Mice used in these studies were obtained from the Jackson Laboratory (C56BL/6J) or Charles River Laboratories (CD-1). Animal procedures performed at University of Alabama at Birmingham (UAB) or Noble Life Sciences (Noble, Woodbine, MD) were conducted in accordance with Public Health Service Policy on the Humane Care and Use of Laboratory Animals and Guide for the Care and Use of Laboratory Animals. Studies performed at UAB were performed under IACUC Approval Number 21203. Studies performed at Noble were performed under the IACUC Approval Number NLS-591.

### Vaccine candidates

Vaccine candidates evaluated in this study were based on a replication-deficient, E1- and E3-deleted adenovirus type 5 vector platform (Tang et al., 2009) and expressed a human codon-optimized gene for either the S1 domain (residues 16 to 685) or the RBD domain (residues 302 to 543) of SARS-CoV-2 spike protein (accession number QHD43416.1). The Ad5-vectored S1 and RBD transgenes included a human tissue plasminogen activator leader sequence and were expressed under the control of the cytomegalovirus immediate early promoter/enhancer. Initial seed stocks were obtained from a transfection of recombinant vector plasmid into E1-complementing PER.C6 cells using a scalable transfection system (Maxcyte STX-100). Cell transfection was performed by a static electroporation using the CL1.1 Processing Assembly procedure (Li et al., 2002). Following vector expansion, replication-deficient vector was obtained from infected cell lysates and was purified over a CsCl gradient, dialyzed against a formulation buffer containing 10 mM Tris at a pH of 7.4, 75 mM NaCl, 1 mM MgCl_2_, 10 mM histidine, 5% (wt/vol) sucrose, 0.02% polysorbate-80 (wt/vol), 0.1 mM EDTA, and 0.5% (vol/vol) ethanol and were then frozen and stored at −65°C.

### Adenovirus vaccine titer measurement

293 HEK cells were seeded in a 96-well plate one day before Ad vector infection. After inoculation of the appropriate dilutions of adenovirus control and test sample(s) onto duplicate wells, the infected cells were incubated for 3 days. At the end of the infection period, media were removed, and cells were fixed with cold methanol. Following drying and rinsing with phosphate-buffered saline, mouse anti-adenovirus-5 hexon antibody was added to each well of cells and incubated at 37°C for at least 60 minutes. After removal of the mouse anti-adenovirus-5 hexon antibody and additional phosphate-buffered saline washes, HRP-conjugated rat anti-mouse antibody was added to each well and incubated at least 60 minutes at 37°C. After removal of the detection antibody and additional phosphate-buffered saline washes, cells were stained with DAB (3, 3-diaminobenzidine) working solution for at least 10 minutes. After removal of DAB working solution and additional washing steps with phosphate-buffered saline, the stained foci were enumerated using a microscope with a 20X objective.

### Vaccination

Inbred female C57BL/6J (The Jackson Laboratory) and outbred CD-1 (Charles River Laboratories) mice of at least 6 weeks of age were randomly allocated into vaccination groups. Replication-deficient Ad5 vector encoding the S1 domain (S1 vector) or the RBD (RBD vector) from the SARS-CoV-2 spike protein were administered intranasally in a volume of 50µl. Dose levels used included high-dose (3.35E+08 ifu), mid-dose (6E+07 ifu) or low-dose (6E+06 ifu) of vaccine except for the S1 vector tested C57BL/6J mice were the high dose vaccine was 6E+08 ifu. The control group received 50µl of buffer alone by intranasal administration.

### Serum Collection

Blood samples were collected from the tail vein of vaccinated mice into BD Microtainer blood collection tubes. The samples were centrifuged at 13000 rpm at room temperature for 8-10 minutes and the serum was collected, aliquoted and frozen at −80°C until analyzed.

### Tissue processing and single cell isolation

Spleen, medLN and lung tissues were isolated from vaccinated mice at the indicated timepoints. Lung tissue was minced and then digested in RPMI-1640 medium containing collagenase (1.25 mg/ml, Sigma) and DNase I (150 units/ml, Sigma) for 45 minutes at 37°C. To generate single cell suspensions, digested lung, spleen, and draining lymph nodes were passed through a fine wire mesh. The single cell suspensions were treated with red blood cell lysis buffer and filtered (100 μm) to remove debris.

### BAL collection

BAL samples were collected using ethyl vinyl acetate (EVA) microbore tubing (Cole-Parmer) with inner and outer diameters of 0.02 in and 0.06 inches, respectively. One end of the tube was fitted to a 23G syringe needle attached to a 3-way stopcock. The other end of the tube was inserted postmortem into an incised trachea. BAL fluid was collected as a single 1ml lavage fraction using ice-cold phenol-free Hank’s Buffered Salt Solution (HBSS, without Ca^2+^ and Mg^2+^) containing 2mM EDTA. BAL cells were separated from the supernatant by centrifugation at 1800 rpm for 5 minutes at 4°C.

### Recombinant SARS-CoV-2 protein production

To produce recombinant SARS-CoV-2 spike ectodomain protein, two human codon-optimized constructs were generated with linear sequence order encoding: a human IgG leader sequence, the SARS-CoV-2 spike ectodomain (amino acids 14-1211), a GGSG linker, T4 fibritin foldon sequence, a GS linker, and finally an AviTag (construct 1) or 6X-HisTag (construct 2). Each construct was engineered with two sets of mutations to stabilize the protein in a pre-fusion conformation. These included substitution of RRAR>SGAG (residues 682 to 685, as in Walls et al., 2020) at the S1/S2 cleavage site and the introduction of two proline residues; K983P, V984P, as in Walls et al., 2020 and Wrapp et al., 2020). Avi/His-tagged trimers were produced by co-transfecting plasmid constructs 1 and 2 (1:2 ratio) into FreeStyle 293-F cells. Cells were grown for three days and the supernatant (media) was recovered by centrifugation. Recombinant spike trimers were purified from media by Fast protein liquid chromatography (FPLC) using a HisTrap HP Column (GE) and elution with 250mM of imidazole. After exchanging into either 10mM Tris-HCl, pH 8.0 or 50mM Bicine, pH 8.3, purified spike ectodomain trimers were biotinylated by addition of biotin-protein ligase (Avidity, Aurora, CO). Biotinylated spike ectodomain trimers were buffer exchanged into PBS, sterile filtered, aliquoted, then stored at −80°C until used.

### SARS-CoV-2 spike cytometric bead array

To generate the spike cytometric bead array (CBA), recombinant SARS-CoV-2 ectodomain trimers were passively absorbed onto streptavidin functionalized fluorescent microparticles (Spherotech 3.6um cat# CPAK-3567-4K, peak 4). 500 µg of biotinylated SARS2-CoV-2 was incubated with 2E+07 Streptavidin functionalized fluorescent microparticles in 400μl of 1% BSA in PBS. Following coupling, the SARS-CoV-2 spike conjugated beads were washed twice in 1ml of 1% BSA, PBS, 0.05% NaN3, resuspended at 1E+08 beads/mL and stored at 4°C. The loading of recombinant SARS2-CoV-2 spike onto the beads was evaluated by staining 1E+05 beads with dilutions ranging from 1μg to 2ng/ml of the recombinant anti-SARS spike antibody CR3022 and visualized with an anti-human IgG secondary antibody.

### CBA IgG and IgA standards

IgG and IgA standards were generated by covalent coupling of isotype-specific polyclonal antibodies to fluorescent particles. Briefly, 0.2 mg of goat polyclonal anti-mouse IgG (southern Biotech cat# 1013-01), anti-IgM (cat#1 022-01), and anti-IgA (cat# 1040-01) antibodies in PBS were mixed with 5E+07 fluorescent microparticles each with a unique fluorescent intensity in the far red channels (Spherotech 3.6um cat# CPAK-3567-4K, peaks 1-3) resuspended in 0.1 M MES buffer pH 5.0. An equal volume of EDC (1-Ethyl-3-(3-dimethylaminopropyl)-carbodiimide), 10mg/mL, in 0.1 M MES (2-(N-morpholino) ethanesulfonic acid) buffer pH 5.0 was added and the mixture was incubated overnight at room temperature. The beads were washed twice by pelleting by centrifugation and resuspension in PBS. Following washing, beads were resuspended in 1% BSA in PBS with 0.005% NaN_3_ as a preservative.

### CBA measurement of spike-specific IgG and IgA responses

The quantification of SARS-CoV-2 spike IgG and IgA was performed in serum or BAL samples obtained from immunized animals using the spike CBA described above. BAL samples (diluted 1/4-8) or serum samples (diluted to 1/1000-5000) in 50µl of PBS were arrayed in 96 well u-bottom polystyrene plates along with 50μl of standards consisting of either mouse IgG, IgM, or IgA ranging from 1µg/ml to 2ng/ml at 0.75x dilutions (Southern Biotech IgM: cat# 0106-01, IgG: cat# 0107-01, IgA cat# 0106-01). 5µl of a suspension containing 5E+05 of each SARS-CoV-2 spike, anti-IgM, anti-IgA, and anti-IgG beads was added to the diluted samples. The suspensions were mixed by pipetting and incubated for 15 minutes at RT. The beads were washed by the addition of 200µl of PBS and centrifuged at 3000g for 5 minutes at RT. The cytometric bead array (CBA) particles were resuspended in a secondary staining solution consisting of polyclonal anti-IgG 488 (southern Biotech cat#1010-30), and either a goat polyclonal anti-IgM (southern Biotech cat# 1020-09) or anti-IgA (southern Biotech cat# 1040-09) conjugated to PE diluted 1/400 in 1% BSA in PBS. The suspension was incubated for 15 minutes in the dark at room temperature. The beads were washed by the addition of 200 µl of PBS and pelleted by centrifugation at 3000g for 5 min at RT. The particles were resuspended in 75µl of PBS and directly analyzed on a BD Cytoflex flow cytometer in plate mode at sample rate of 100 μl per minute. Sample collection was stopped following the acquisition of 75 µL. Following acquisition, the resulting FCS files were analyzed in flowJo (treestar). Briefly, the beads were identified by gating on singlet 3.6 μm particles in log scale in the forward scatter and side scatter parameters. APC-Cy7 channel fluorescence gates were used to segregate the particles by bead identity. Geometric mean fluorescent intensity was calculated in the PE and 488 channels. Best fit power curves were generated from the Ig capture beads using the known concentration of standards on a plate by plate basis. This formula was applied to the MFI of the SARS-COV-2 spike particles for all samples of the corresponding assay converting MFI to ng/ml or µg/ml. These calculated values were corrected for the dilution factor.

### Neutralizing antibody titers

A focus reduction neutralization test (FRNT) was used to quantify the titer of neutralizing antibodies against SARS-CoV-2 isolate USA-WA1/2020. Vero E6 cells were grown on 96-well plates to confluence. On the day of the infection phase of the assay, serial dilutions (1:20-1:2560) of antisera were made and combined and incubated with an equal volume of viral stock, at a specified dilution for 30 minutes at room temperature, such that the final dilutions of antisera ranged from 1:40 to 1:5120. The viral stock was diluted from a concentrated working stock to produce an estimated 30 viral focal units per well. After incubation, the sera:virus mixtures were added to the wells (100µL), and infection allowed to proceed for 1 hour on the Vero cells at 35°C. At the completion of the 1-hour incubation, a viscous overlay of Eagle’s MEM with 4% FBS and antibiotics and 1.2% Avicell were added to sera:virus mixture on the cell monolayers such that the final volume was 200 µL per well. The infection was allowed to proceed for 24 hr. The next day, each plate was fixed by submerging the entire plate and contents in 10% formalin/PBS for 24 h. Detection of virus foci reduction was performed on the fixed 96 well plates. Briefly, plates were rinsed in H_2_O, and methanol:hydrogen peroxide added to the wells for 30 min with rocking to quench endogenous peroxidase activity. After quenching, plates were rinsed in H_2_O to remove methanol and 5% Blotto was added to the wells as a blocking solution for 1 hour. For primary antibody detection, a SARS-CoV-2 spike/RBD antibody (Rabbit, Polyclonal, SinoBiologicals Ct No. 40592-T62) was added to 5% Blotto and incubated on the monolayers overnight. Plates were washed 5X with PBS, and further incubated with a goat anti-rabbit IgG conjugated to horseradish peroxidase (Boster Biological Technology Co., #BA1054-0.5) in 5% Blotto for 1 hr. Plates were rinsed once with 0.05% Tween in 1X PBS followed by 5 washes in 1X PBS. Impact DAB detection kit (Vector Labs #SK-4105) was used to detect peroxidase activity. Brown foci were counted manually from the scanned image of each well, recorded, and the reduction of foci as compared to equivalent naïve mouse sera controls was determined. FRNT_50_ titers were calculated using a 4PL curve fit to determine the serum dilution corresponding to a 50% reduction in the foci present in control wells.

### Flow cytometry analysis of innate and adaptive immune cells

Cell numbers per tissue were determined by mixing 20 µl of each single cell suspension into a 96-well plate with 50 μl of 8.4 × 10^4^ Fluoresbrite Carboxylate YG 10 µm microspheres/ml (Polysciences Inc.) and 180 μl staining media (dPBS + 2% FBS) containing 2 mM EDTA (SME) and 7-AAD (1:720 dilution). To perform flow cytometric analysis, 200 µl of each sample were placed into 3 separate V-bottom 96-well plates for antibody staining for flow cytometric analysis. Samples were incubated for 10 minutes at 4°C in the dark with Fc-Block (clone 24G2, 10 μg/ml), washed with 200 µl staining media (SME) and then stained with myeloid, B cell or BAL antibody panels. The myeloid panel consisted of B220/CD45R-FITC (clone RA3-6B2; 1:200 dilution), Ly6G-PE (clone 1A8; 1:200 dilution), CD64-PerCP-Cy5.5 (clone X54-5/7.1; 1:150 dilution), CD11b-APC (clone M1/70; 1:200 dilution), CD11c-PE-Cy7 (clone N418; 1:300 dilution), Ly6C-APC-Cy7 (clone AL-21; 1:200 dilution), MHCII-PB (clone M5/114.15.4; 1:600 dilution) and Aqua LIVE/DEAD (1:1000 dilution). Cells stained with the myeloid panel were incubated with the antibody mix (50 μl total volume) for 20 minutes at 4°C in the dark. The B cell panel consisted of CD95/FAS-FITC (clone Jo2; 1:200 dilution), F4/80-PerCP-Cy5.5 (clone BM8; 1:200 dilution), CD3-PerCP-Cy5.5 (clone 17A2; 1:200 dilution), CD38-PE-Cy7 (clone 90; 1:400 dilution), CD19-APC-Fire750 (clone 6D5; 1:200 dilution), CD138-BV421 (clone 281-2; 1:200 dilution) and IgD-BV510 (clone 11-26c.2a; 1:500 dilution). Cells stained with the B cell panel were incubated with antibody mix (50 μl total volume) for 45 min at 4°C in the dark. The BAL panel consisted of Ly6G-PE (clone 1A8; 1:200 dilution), CD64-PerCP-Cy5.5 (clone X54-5/7.1; 1:150 dilution), CD8a-APC (clone 53-6.7; 1:200 dilution), CD11c-PE-Cy7 (clone N418; 1:200 dilution), CD19-APC/Fire750 (clone 6D5; 1:200 dilution; CD4-eFLUOR450 (clone GK1.5; 1:200 dilution) and Aqua LIVE/DEAD (1:1000 dilution). Cells stained with the BAL panel were incubated with the antibody mix (50 μl total volume) for 20 min at 4°C in the dark. Following incubation with the different flow cytometry panels, the cells were washed, resuspended in 100 µl of eBioscience fixation buffer (00-5521-00) and incubated for 20 min at 4 °C. After incubation, the cells were spun down and resuspended in 200 μl SME. Stained and fixed cells from all antibody panels and cell counting panels were analyzed on a BD Canto II.

### Synthetic RBD peptides

For analysis of T cell responses, a pool of 53 peptides derived from a peptide scan through RBD of spike protein of SARS-CoV-2 (319-541) was designed and synthesized by JPT (JPT Peptide technologies, Berlin, Germany). Peptides were designed with a length of 15 a.a. and an overlap of 11a.a. Before use, each vial containing 15 nmol (appr. 25µg) of each peptide per vial was reconstituted in 50µl of DMSO before dilution into complete culture media.

### IFN-γ ELIspot

Spleen and lung cell suspensions (150,000 cells/well) were placed in individual wells of ELIspot plates (Millipore-Sigma) that were pre-coated with anti-IFN-γ (AN18, 5 μg/ml). Cells were stimulated with the RBD peptide pool described above at 2.0 μg/peptide/ml. Following 24 hr stimulation, plates were stained with biotinylated anti-IFN-γ (R4-6A2), followed by washing steps, and incubation with streptavidin-ALP. Secreted IFN-γ was detected following incubation with NBT/NCPI substrate for 7-10 min. The number of IFN-γ spot-forming cells were manually counted from digital images of each well.

### Intracellular cytokine staining

The analysis of CD4+ and CD8+ T cell responses in lung tissues and spleens by flow cytometry was performed as follows. Spleen and lung single cell suspensions were stimulated with the RBD peptide pool for 5 hrs in the presence of Brefeldin A (5 hrs, 12.5 ug/mL concentration). Cells were then incubated on ice with a combination of fluorescent dye-labelled antibodies including anti-CD4-V500 (clone GK1.5; 1:200 dilution), anti-CD8α-APC-Fire750 (clone 53-6.7; 1:200 dilution), anti-CD11a/CD18-Pacific Blue (H155-78; 1:200 dilution), anti-CD103-PE (M290; 1:200 dilution), anti-CD69-FITC (H1-2F3; 1:200 dilution), anti-Ly6G-PerCP-Cy5.5 (clone 1A8; 1:200 dilution), anti-CD64-PerCP-Cy5.5 (clone X54-5/7.1; 1:200 dilution), anti-B220/CD45R-PerCP (clone RA3-6B2; 1:200 dilution), and Red LIVE/DEAD (1:1000 dilution). Following surface staining, cells were permeabilized using BD Biosciences Cytofix/Cytoperm kit and stained with anti-IFN-γ-PE-Cy7 (XMG1.2; 1:200 dilution) and anti-TNF-α-APC (MP6-XT22; 1:200 dilution). Following incubation with the antibodies, cells were washed and resuspended before analysis on FACSCanto II within 12 hours.

### Measurement of inflammatory cytokines in culture supernatants

Protein levels of IFNγ, IL-2, IL-4, IL-5, IL-10, IL-13, IL-17A and TNFα were quantified in culture supernatants using the mouse-specific Milliplex® multi-analyte panel kit MT17MAG-47K (Millipore; Sigma) and the MagPix® instrument platform with related xPONENT® software (Luminex Corporation). The readouts were analyzed with the standard version of EMD Millipore’s Milliplex® Analyst software.

### Quantification and statistical analysis

Statistical significance was assigned when P values were < 0.05 using Prism Version 8.4.2 (GraphPad). All tests and values are indicated in the relevant Figure legends.

## Supplementary Figures

**Supplementary Figure 1:**
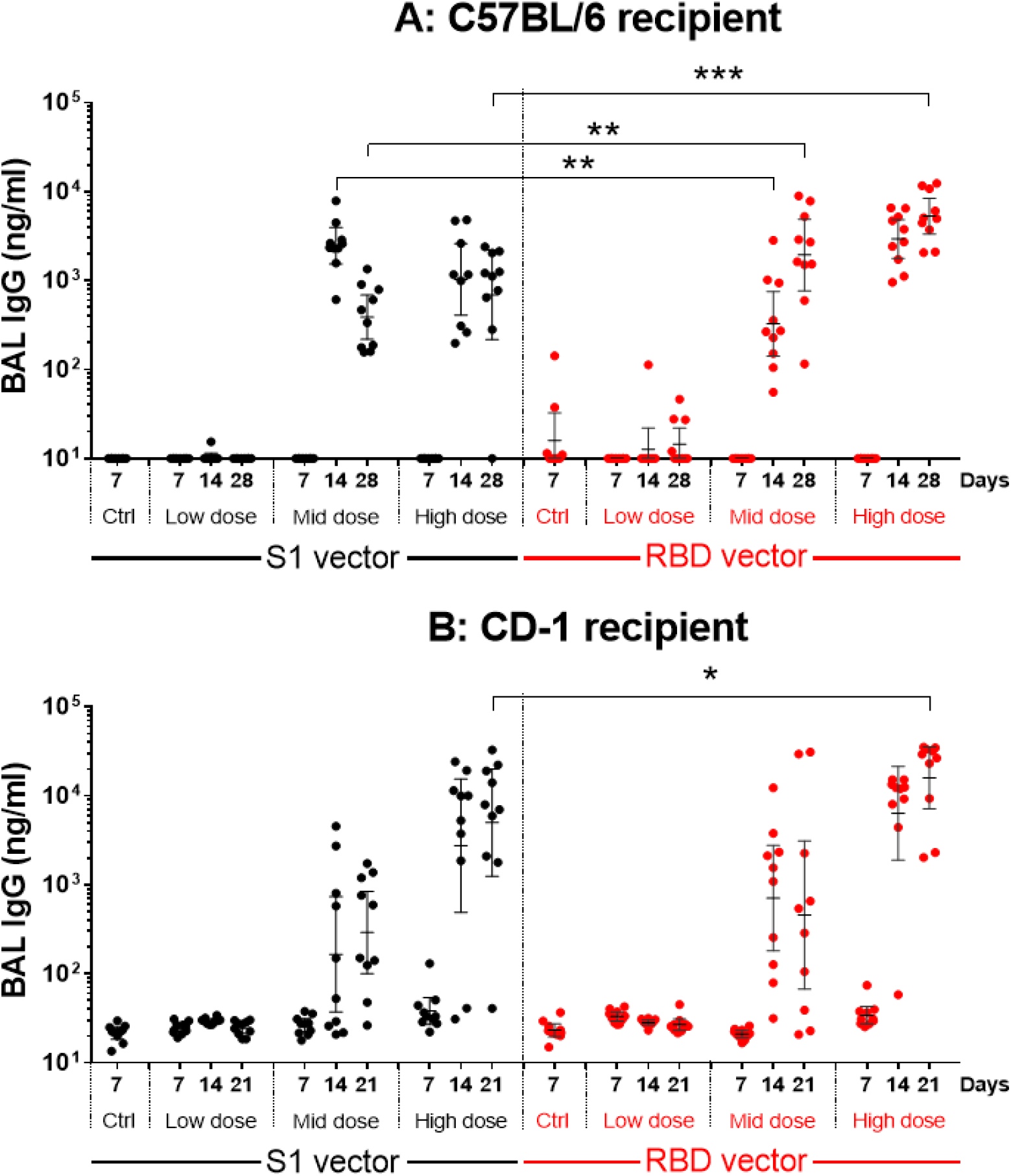
Spike-specific IgG responses in BALs following single intranasal administration of S1 and RBD vectors. C57BL/6J (A) and CD-1 (B) mice received a single intranasal administration of vehicle (Ctrl), the S1 Ad5 vaccine (S1 vector) or RBD Ad5 vaccine (RBD vector) given at a low, middle (mid) or high dose as described in Material and Methods. BAL samples were collected at the indicated timepoints (n=10 animals/group/timepoint) and analyzed individually for the quantification of Spike-specific IgG. Results are expressed in ng/ml. Lines represent geometric mean response +/− 95% confidence interval. Statistical analysis was performed with a Mann-Whitney test: *, P < 0.05; **, P <0.01; ***, P < 0.001; ****, P < 0.0001.

**Supplementary Figure 2:**
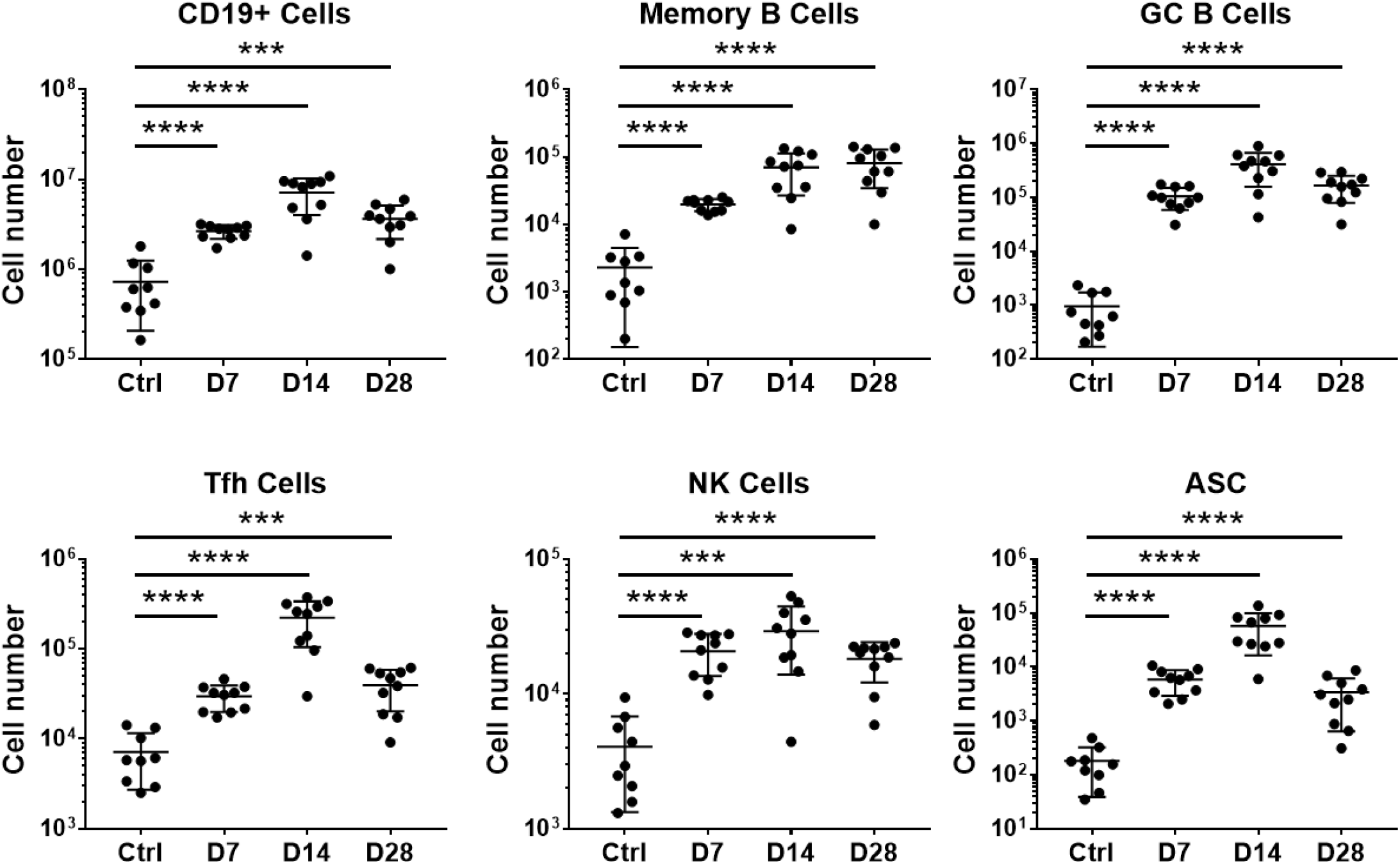
Flow cytometry analysis of immune cells in mediastinal lymph nodes from C57BL/6J mice following intranasal vaccination with a single high dose of the RBD vector (AdCOVID). C57BL/6J mice were given a single intranasal administration of vehicle (Ctrl) or the high dose RBD vector (AdCOVID) as described in Material and Methods. Mediastinal lymph node cells were isolated from vaccinated animals at the indicated timepoints (10 mice/timepoint) and analyzed individually by flow cytometry as described in Materials and Methods. Results are expressed as cell number. Lines represent mean response +/− SD. Statistical analysis was performed with Mann-Whitney test: *, P < 0.05; **, P <0.01; ***, P < 0.001; ****, P < 0.0001.

**Supplementary Figure 3:**
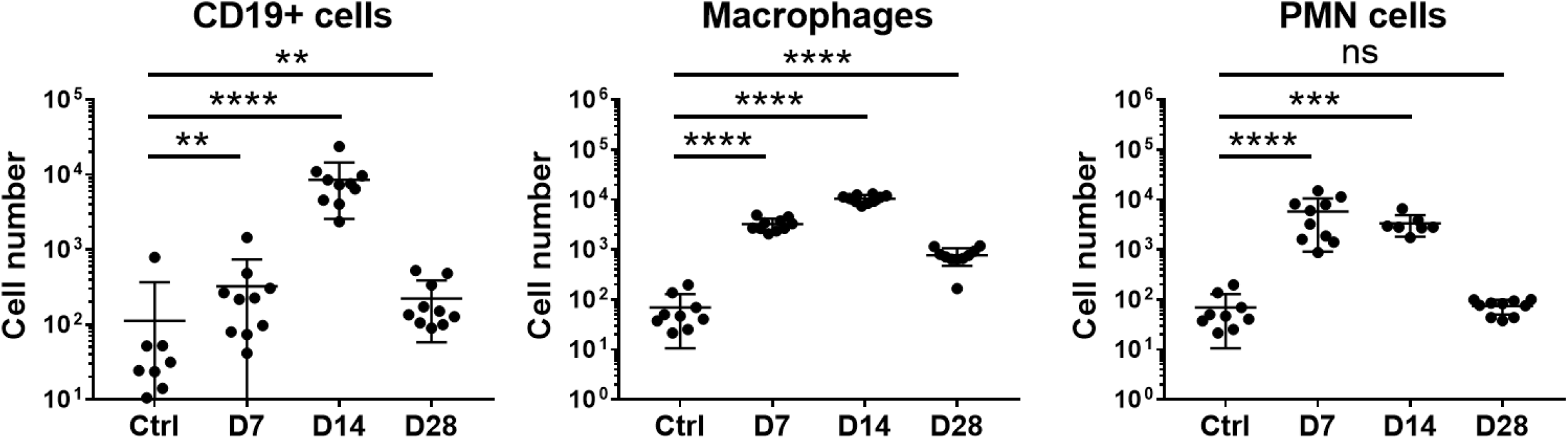
Flow cytometry analysis of immune cells in BAL from C57BL/6J mice following intranasal vaccination with a single high dose of RBD vector (AdCOVID). C57BL/6J mice were given a single intranasal administration of vehicle (Ctrl) or the high dose RBD vector (AdCOVID) as described in Material and Methods. BAL cells were collected from vaccinated animals at the indicated timepoints (10 mice/timepoint) and analyzed individually by flow cytometry as described in Materials and Methods. Results are expressed as cell number. Lines represent mean response +/− SD. Statistical analysis was performed with a Mann-Whitney test: *, P < 0.05; **, P <0.01; ***, P < 0.001; ****, P < 0.0001.

**Supplementary Figure 4:**
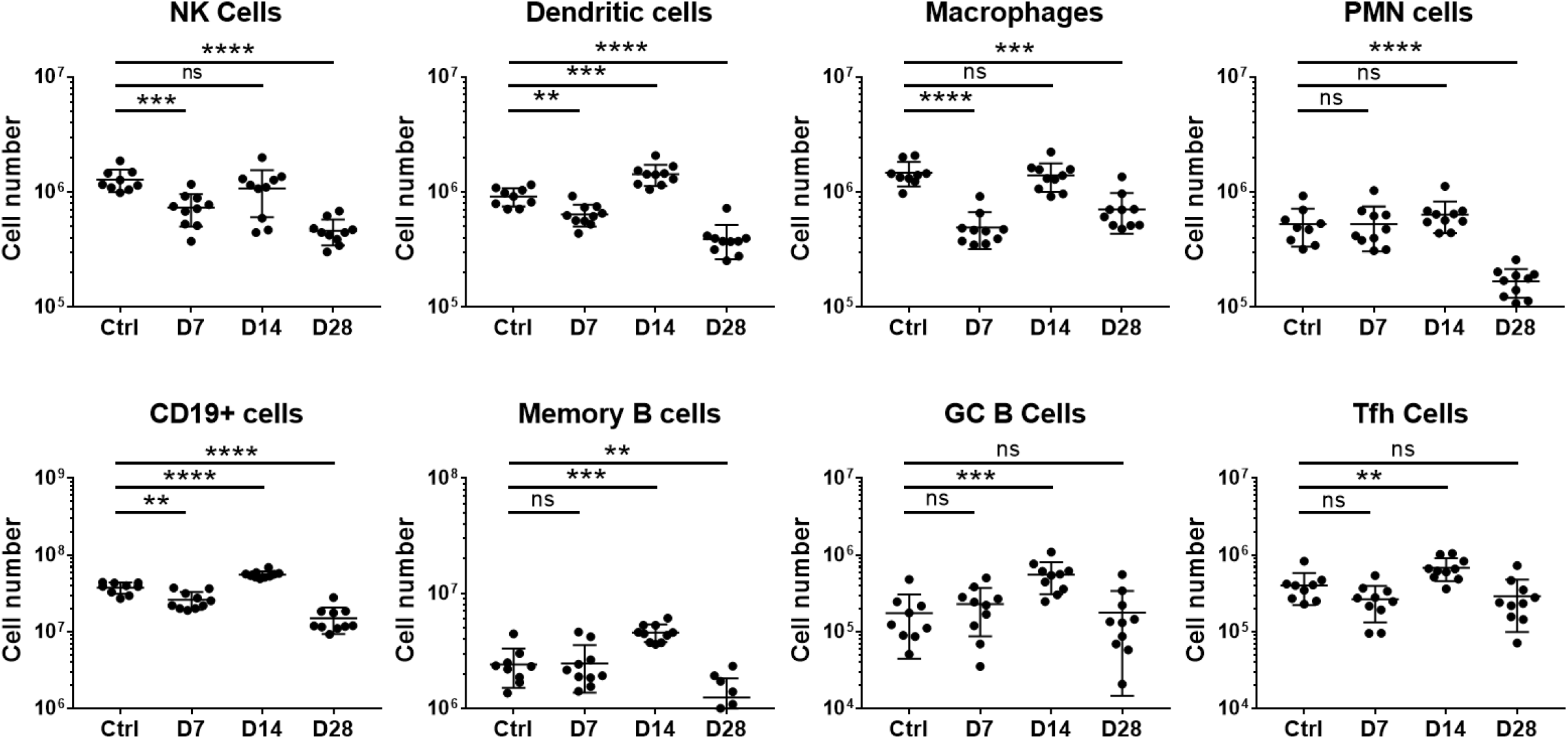
Flow cytometry analysis of immune cells in spleens from C57BL/6J mice following intranasal vaccination with a single high dose of RBD vector (AdCOVID). C57BL/6J mice were given a single intranasal administration of vehicle (Ctrl) or the high dose RBD vector (AdCOVID) as described in Material and Methods. Spleen cells isolated collected from vaccinated animals at the indicated timepoints (10 mice/timepoint) and analyzed individually by flow cytometry as described in Materials and Methods. Results are expressed as cell number. Lines represent mean response +/− SD. Statistical analysis was performed with Mann-Whitney test: *, P < 0.05; **, P <0.01; ***, P < 0.001; ****, P < 0.0001

## Notes

### Competing Interest Statement

The authors have declared no competing interest.

## References

Advisory Board: What we know (so far) about the long-term health effects of COVID-19 (2020). https://www.advisory.com/daily-briefing/2020/06/02/covid-health-effects.

Belshe RB, Gruber WC, Mendelman PM, Mehta HB, Mahmood K, Reisinger K, Treanor J, Zangwill K, Hayden FG, Bernstein DI, Kotloff K, King J, Piedra PA, Block SL, Yan L, Wolff M. Correlates of immune protection induced by live, attenuated, cold-adapted, trivalent, intranasal influenza virus vaccine. J Infect Dis. 2000 Mar;181(3):1133–7.

Bissett C, Gilbride C, Williamson BN, Rosenke R, Long D, Ishwarbhai A, Kailath R, Rose L, Morris S, Powers C, Lovaglio J, Hanley PW, Scott D, Saturday G, de Wit E, Gilbert SC, Munster VJ. ChAdOx1 nCoV-19 vaccine prevents SARS-CoV-2 pneumonia in rhesus macaques. Nature. 2020 Jul 30.

Boyaka PN. Inducing Mucosal IgA: A Challenge for Vaccine Adjuvants and Delivery Systems. J Immunol. 2017 Jul 1;199(1):9–16.

Brann DH, Tsukahara T, Weinreb C, Lipovsek M, Van den Berge K, Gong B, Chance R, Macaulay IC, Chou HJ, Fletcher RB, Das D, Street K, de Bezieux HR, Choi YG, Risso D, Dudoit S, Purdom E, Mill J, Hachem RA, Matsunami H, Logan DW, Goldstein BJ, Grubb MS, Ngai J, Datta SR. Non-neuronal expression of SARS-CoV-2 entry genes in the olfactory system suggests mechanisms underlying COVID-19-associated anosmia. Sci Adv. 2020 Jul 31;6(31):eabc5801.

Centers for Disease Control and Prevention. Coronavirus disease 2019 (COVID-19): People who are at increased risk for severe illness (2020a). Available at: https://www.cdc.gov/coronavirus/2019-ncov/need-extra-precautions/people-at-increased-risk.html. Accessed 4 August 2020.

Centers for Disease Control and Prevention. Coronavirus disease 2019 (COVID-19): Scientific Brief: SARS-CoV-2 and Potential Airborne Transmission (2020b). Available at: https://www.cdc.gov/coronavirus/2019-ncov/more/scientific-brief-sars-cov-2.html. Accessed 5 October 2020.

Croyle MA, Patel A, Tran KN, et al. Nasal delivery of an adenovirus-based vaccine bypasses pre-existing immunity to the vaccine carrier and improves the immune response in mice. PLoS One. 2008;3(10):e3548.

Fransen F, Zagato E, Mazzini E, Fosso B, Manzari C, El Aidy S, Chiavelli A, D’Erchia AM, Sethi MK, Pabst O, Marzano M, Moretti S, Romani L, Penna G, Pesole G, Rescigno M. BALB/c and C57BL/6 Mice Differ in Polyreactive IgA Abundance, which Impacts the Generation of Antigen-Specific IgA and Microbiota Diversity. Immunity. 2015 Sep 15;43(3):527–40.

Funk CD, Laferrière C, Ardakani A. A Snapshot of the Global Race for Vaccines Targeting SARS-CoV-2 and the COVID-19 Pandemic. Front Pharmacol. 2020;11:937.

Gould VMW, Francis JN, Anderson KJ, Georges B, Cope AV, Tregoning JS. Nasal IgA Provides Protection against Human Influenza Challenge in Volunteers with Low Serum Influenza Antibody Titre. Front Microbiol. 2017 May 17;8:900.

Guo YR, Cao QD, Hong ZS, Tan YY, Chen SD, Jin HJ, Tan KS, Wang DY, Yan Y. The origin, transmission and clinical therapies on coronavirus disease 2019 (COVID-19) outbreak – an update on the status. Mil Med Res. 2020 Mar 13;7(1):11.

Hassan AO, Kafai NM, Dmitriev IP, Fox JM, Smith BK, Harvey IB, Chen RE, Winkler ES, Wessel AW, Case JB, Kashentseva E, McCune BT, Bailey AL, Zhao H, VanBlargan LA, Dai YN, Ma M, Adams LJ, Shrihari S, Danis JE, Gralinski LE, Hou YJ, Schäfer A, Kim AS, Keeler SP, Weiskopf D, Baric RS, Holtzman MJ, Fremont DH, Curiel DT, Diamond MS. A Single-Dose Intranasal ChAd Vaccine Protects Upper and Lower Respiratory Tracts against SARS-CoV-2. Cell. 2020 Oct 1;183(1):169–184.e13.

Hoffmann M, Kleine-Weber H, Schroeder S, et al. SARS-CoV-2 cell entry depends on ACE2 and TMPRSS2 and is blocked by a clinically proven protease inhibitor. Cell. 2020;181(2):271–280.e8.

Holmgren J, Czerkinsky C. Mucosal immunity and vaccines. Nat Med. 2005 Apr;11(4 Suppl):S45–53.

Hou YJ, Okuda K, Edwards CE, Martinez DR, Asakura T, Dinnon KH 3rd, Kato T, Lee RE, Yount BL, Mascenik TM, Chen G, Olivier KN, Ghio A, Tse LV, Leist SR, Gralinski LE, Schäfer A, Dang H, Gilmore R, Nakano S, Sun L, Fulcher ML, Livraghi-Butrico A, Nicely NI, Cameron M, Cameron C, Kelvin DJ, de Silva A, Margolis DM, Markmann A, Bartelt L, Zumwalt R, Martinez FJ, Salvatore SP, Borczuk A, Tata PR, Sontake V, Kimple A, Jaspers I, O’Neal WK, Randell SH, Boucher RC, Baric RS. SARS-CoV-2 Reverse Genetics Reveals a Variable Infection Gradient in the Respiratory Tract. Cell. 2020 Jul 23;182(2):429–446.e14.

Krishnan V, Andersen BH, Shoemaker C, et al. Efficacy and immunogenicity of single-dose AdVAV intranasal anthrax vaccine compared to anthrax vaccine absorbed in an aerosolized spore rabbit challenge model. Clin Vaccine Immunol. 2015;22(4):430–439.

Laidlaw BJ, Zhang N, Marshall HD, Staron MM, Guan T, Hu Y, Cauley LS, Craft J, Kaech SM. CD4+ T cell help guides formation of CD103+ lung-resident memory CD8+ T cells during influenza viral infection. Immunity. 2014 Oct 16;41(4):633–45.

Letko M, Marzi A, Munster V. Functional assessment of cell entry and receptor usage for SARS-CoV-2 and other lineage B betacoronaviruses. Nature Microbiology 2020;5: 562–569.

Li LH, Shivakumar R, Feller S, Allen C, Weiss JM, Dzekunov S, Singh V, Holaday J, Fratantoni J, Liu LN. Highly efficient, large volume flow electroporation. Technol Cancer Res Treat. 2002 Oct;1(5):341–50.

Li H, Wang Y, Ji M, Pei F, Zhao Q, Zhou Y, Hong Y, Han S, Wang J, Wang Q, Li Q, Wang Y. Transmission Routes Analysis of SARS-CoV-2: A Systematic Review and Case Report. Front Cell Dev Biol. 2020 Jul 10;8:618.

Lowen AC, Steel J, Mubareka S, Carnero E, García-Sastre A, Palese P. Blocking interhost transmission of influenza virus by vaccination in the guinea pig model. J Virol. 2009 Apr;83(7):2803–18. doi: 10.1128/JVI.02424-08. Epub 2009 Jan 19.

Pizzolla A, Nguyen THO, Smith JM, Brooks AG, Kedzieska K, Heath WR, Reading PC, Wakim LM. Resident memory CD8+ T cells in the upper respiratory tract prevent pulmonary influenza virus infection. Sci Immunol. 2017 Jun 2;2(12):eaam6970.

Premkumar L, Segovia-Chumbez B, Jadi R, Martinez DR, Raut R, Markmann A, Cornaby C, Bartelt L, Weiss S, Park Y, Edwards CE, Weimer E, Scherer EM, Rouphael N, Edupuganti S, Weiskopf D, Tse LV, Hou YJ, Margolis D, Sette A, Collins MH, Schmitz J, Baric RS, de Silva AM. The receptor binding domain of the viral spike protein is an immunodominant and highly specific target of antibodies in SARS-CoV-2 patients. Sci Immunol. 2020 Jun 11;5(48):eabc8413.

Price GE, Lo CY, Misplon JA, Epstein SL. Mucosal immunization with a candidate universal influenza vaccine reduces virus transmission in a mouse model. J Virol. 2014 Jun;88(11):6019–30.

Ravichandran S, Coyle EM, Klenow L, Tang J, Grubbs G, Liu S, Wang T, Golding H, Khurana S. Antibody signature induced by SARS-CoV-2 spike protein immunogens in rabbits. Sci Transl Med. 2020 Jul 1;12(550):eabc3539.

Sekine T, Perez-Potti A, Rivera-Ballesteros O, et al. Robust T cell immunity in convalescent individuals with asymptomatic or mild COVID-19 [published online ahead of print, 2020 Aug 14]. Cell. 2020;doi:10.1016/j.cell.2020.08.017.

Seydoux E, Homad LJ, MacCamy AJ, et al. Characterization of neutralizing antibodies from a SARS-CoV-2 infected individual. Preprint. bioRxiv. 2020;2020.05.12.091298.

Sungnak W, Huang N, Bécavin C, et al. SARS-CoV-2 entry factors are highly expressed in nasal epithelial cells together with innate immune genes. Nat Med. 2020;26(5):681–687.

Takamura S. Persistence in Temporary Lung Niches: A Survival Strategy of Lung-Resident Memory CD8+ T Cells. Viral Immunol. 2017 Jul/Aug;30(6):438–450.

Tang DC, Zhang J, Toro H, Shi Z, Van Kampen KR. Adenovirus as a carrier for the development of influenza virus-free avian influenza vaccines. Expert Rev Vaccines. 2009 Apr;8(4):469–81.

van Doremalen N, Lambe T, Spencer A, Belij-Rammerstorfer S, Purushotham JN, Port JR, Avanzato VA, Bushmaker T, Flaxman A, Ulaszewska M, Feldmann F, Allen ER, Sharpe H, Schulz J, Holbrook M, Okumura A, Meade-White K, Pérez-Pérez L, Edwards NJ, Wright D, Bissett C, Gilbride C, Williamson BN, Rosenke R, Long D, Ishwarbhai A, Kailath R, Rose L, Morris S, Powers C, Lovaglio J, Hanley PW, Scott D, Saturday G, de Wit E, Gilbert SC, Munster VJ. ChAdOx1 nCoV-19 vaccine prevents SARS-CoV-2 pneumonia in rhesus macaques. Nature. 2020 Jul 30.

van Ginkel FW, Nguyen HH, McGhee JR. Vaccines for mucosal immunity to combat emerging infectious diseases. Emerg Infect Dis. 2000 Mar-Apr;6(2):123–32.

Walls AC, Park YJ, Tortorici MA, Wall A, McGuire AT, Veesler D. Structure, Function, and Antigenicity of the SARS-CoV-2 Spike Glycoprotein. Cell. 2020;181(2):281–292 e286.

Walsh EE, Frenck R, Falsey AR, Kitchin N, Absalon J, Gurtman A, Lockhart S, Neuzil K, Mulligan MJ, Bailey R, Swanson KA, Li P, Koury K, Kalina W, Cooper D, Fontes-Garfias C, Shi PY, Türeci Ö, Thompkins KR, Lyke KE, Raabe V, Dormitzer PR, Jansen KU, Sahin U, Gruber WC. RNA-Based COVID-19 Vaccine BNT162b2 Selected for a Pivotal Efficacy Study. medRxiv [Preprint]. 2020 Aug 20:2020.08.17.20176651.

World Health Organization. Coronavirus disease 2019 (COVID-19) (2020a). https://covid19.who.int/. Accessed on October 05,2020.

World Health Organization. Draft landscape of COVID-19 candidate vaccines (2020b). Available at: https://www.who.int/publications/m/item/draft-landscape-of-covid-19-candidate-vaccines. Accessed 2 October 2020

Wrapp D, Wang N, Corbett KS, et al. Cryo-EM structure of the 2019-nCoV spike in the prefusion conformation. Science. 2020;367(6483):1260–1263.

Wu S, Zhong G, Zhang J, Shuai L, Zhang Z, Wen Z, Wang B, Zhao Z, Song X, Chen Y, Liu R, Fu L, Zhang J, Guo Q, Wang C, Yang Y, Fang T, Lv P, Wang J, Xu J, Li J, Yu C, Hou L, Bu Z, Chen W. A single dose of an adenovirus-vectored vaccine provides protection against SARS-CoV-2 challenge. Nat Commun. 2020 Aug 14;11(1):4081.

Yusuf H, Kett V. Current prospects and future challenges for nasal vaccine delivery. Hum Vaccin Immunother. 2017;13(1):34–45.

Zhang J, Tarbet EB, Feng T, Shi Z, Van Kampen KR, Tang DC. Adenovirus-vectored drug-vaccine duo as a rapid-response tool for conferring seamless protection against influenza. PLoS One. 2011;6(7):e22605.

Zhu N, Zhang D, Wang W, Li X, Yang B, Song J, Zhao X, Huang B, Shi W, Lu R. et al. A novel coronavirus from patients with pneumonia in China, 2019. N Engl J Med. 2020;382:727–733.

